# Lis1 mutation prevents basal radial glia-like cell production in the mouse

**DOI:** 10.1101/2020.10.10.334508

**Authors:** Maxime Penisson, Shinji Hirotsune, Fiona Francis, Richard Belvindrah

**Affiliations:** INSERM U 1270, Paris, France; Sorbonne University, UMR-S 1270, F-75005 Paris, France; Institut du Fer à Moulin, Paris, France; Osaka University, Osaka Japan

**Author notes:** These authors contributed equally (Fiona Francis, Richard Belvindrah). Corresponding author: Fiona Francis, Institut du Fer à Moulin, 17 rue du Fer à Moulin, 75005, Paris France.

## Abstract

Human cortical malformations are associated with progenitor proliferation and neuronal migration abnormalities. Progenitor cells include apical radial glia, intermediate progenitors and basal (or outer) radial glia (bRGs or oRGs). bRGs are few in number in lissencephalic species (e.g. the mouse) but abundant in gyrencephalic brains. The *LIS1* gene coding for a dynein regulator, is mutated in human lissencephaly, associated also in some cases with microcephaly. LIS1 was shown to be important during cell division and neuronal migration. Here, we generated bRG-like cells in the mouse embryonic brain, investigating the role of Lis1 in their formation. This was achieved by *in utero* electroporation of a hominoid-specific gene *TBC1D3* (coding for a RAB-GAP protein) at mouse embryonic day (E) 14.5. We first confirmed that *TBC1D3* expression in wild-type (WT) brain generates numerous Pax6^+^ bRG-like cells that are basally localized. Second, using the same approach, we assessed the formation of these cells in heterozygote *Lis*1 mutant brains. Our novel results show that *Lis*1 depletion in the forebrain from E9.5 prevented subsequent *TBC1D*3-induced bRG-like cell amplification. Indeed, we observe disruption of the ventricular zone (VZ) in the mutant. *Lis*1 depletion altered adhesion proteins and mitotic spindle orientations at the ventricular surface and increased the proportion of abventricular mitoses. Progenitor outcome could not be further altered by TBC1D3. We conclude that perturbation of *Lis*1/LIS1 dosage is likely to be detrimental for appropriate progenitor number and position, contributing to lissencephaly pathogenesis.

## Introduction

The development of the cerebral cortex relies on different types of progenitor cell situated in a neuroepithelium adjacent to the cerebral ventricles. These cells produce neurons which *in fine* will form networks that underlie brain functions (Taverna et al., 2014). Early neuroepithelial cells give rise to apical radial glia cells (aRGs) which are localized in the ventricular zone (VZ) and possess a short apical process descending to the ventricle and a long basal process extending to the pial surface (Götz and Huttner, 2005). These progenitor cells proliferate producing either immature glutamatergic neurons which will migrate along aRG basal fibers in the intermediate zone (IZ) to reach their final position in the cortical plate (CP), or other subpopulations of progenitors: e.g. intermediate progenitors (IPs) and basal radial glial cells (bRGs). IPs are multipolar cells that are localized mainly in the subventricular zone (SVZ) and can produce deep and upper layer neurons (Agirman et al., 2017; Hevner, 2019). bRGs are rare in the rodent, but abundant in gyrencephalic species where they are localized in an outer SVZ (oSVZ). In these species, they have been shown to play a major role during cortical development (Florio et al., 2015; Hansen et al., 2010; Fietz et al., 2010; Penisson et al., 2019; Pollen et al., 2015; Reillo et al., 2011). They are highly proliferative cells and can produce neurons and IPs, but unlike rodent IPs, they have the ability to self-renew extensively, therefore constituting a renewed pool of progenitors for cortical development (Betizeau et al., 2013; Hansen et al., 2010; LaMonica et al., 2012). Basal RGs present different morphologies, classically they were described with only a basal process, but they can also have only an apical process or both (Betizeau et al., 2013). It is important to assess how perturbation of bRG function may contribute to the apparition of cortical disorders.

Malformations of cortical development (MCDs) are rare pathologies characterized by intellectual disability and/or epilepsy. They are associated with abnormalities in cortical structure and/or the number of neurons (Desikan and Barkovich, 2016). Linked to genetic mutations or environmental factors (Romero et al., 2018), they have been associated with defects in cell proliferation and neuronal migration, and include micro- or macrocephaly (reduction or enlargement of cerebral volume respectively), lissencephaly (absence of or abnormal folds) or heterotopias (presence of grey matter within the white matter) (Barkovich et al., 2012; Capuano et al., 2017; Subramanian et al., 2020).

*LIS1*, coding for a regulator of dynein activity, was the first gene to be linked with a neuronal migration disorder: the Miller-Dieker syndrome (MDS) (Dobyns et al., 1993; Reiner and Sapir, 2013, Cianfrocco et al., 2015). MDS is characterized by lissencephaly and facial abnormalities and is caused by a contiguous deletion of genes on the short arm of chromosome 17. While several genes are deleted in MDS, *LIS*1 appears to be one of the main actors in this pathology, as heterozygous intragenic deletions and point mutations also lead to lissencephaly, with varying degrees of severity, including microcephaly (Lo Nigro et al., 1997; Yingling et al., 2003).

In the naturally lissencephalic mouse, *Lis1* haploinsufficiency leads to mild neocortical and hippocampal disorganization, contrasting with the extremely severe human disorder. Only further depletion of *Lis1* to approximately 35 % of normal, induced a severe neocortical phenotype (Gambello et al., 2003; Hirotsune et al., 1998). Reduced *Lis1* dosage was shown to affect neuronal migration (Tsai et al., 2005), as well as mitotic spindle orientation in neuroepithelial cells and aRGs (Yingling et al., 2008). This leads to a depletion of the progenitor pool, by increased cell cycle exit of aRGs transiently favorising neurogenesis (Iefremova et al., 2017; Pramparo et al., 2010). Human MDS organoids were produced (Bershteyn et al., 2017; Iefremova et al., 2017) revealing, as well as other defects, bRGs with longer mitoses and altered mitotic somal translocation (Bershteyn et al., 2017). However, the specific role of LIS1 in bRG generation still remains poorly understood, as well as more generally the involvement of bRGs in the pathogenesis of lissencephaly. There is indeed a strong need to revisit MCD pathogenesis, considering the more recently identified bRGs, questioning how perturbation of their function might contribute to these disorders.

Various genes and mechanisms have been described during the past decade as regulating bRG production and amplification (Penisson et al., 2019). Several models exist to study these cells in different organisms. Certain reports have described an artificial enrichment of bRG-like cells in the mouse brain (Florio et al., 2015; Heng et al., 2017; Ju et al., 2016; Liu et al., 2017; Lui et al., 2014; Stahl et al., 2013; Tavano et al., 2018; Wong et al., 2015), sometimes even leading to the formation of folds on the surface of the brain. Hence, with the aim of studying the role of Lis1 in the genesis and function of bRGs, we set out to combine the study of a floxed mouse line for *Lis1* (Hirotsune et al., 1998) with an amplification of bRG-like cells. We selected to use TBC1D3, a hominoid-specific RAB-GAP, known to favorize the generation of bRG-like cells upon expression in the mouse brain (Ju et al., 2016).

In this study, we confirmed that *TBC1D3* expression in the mouse via *in utero* electroporation (IUE) at E14.5 promotes the generation of bRG-like cells 2 days later. We found that an early heterozygote depletion of *Lis1* at E9.5 using the Emx1-Cre mouse line (Gorski et al., 2002) prevents the TBC1D3-dependent bRG-like cell amplification which occurs in control animals. Indeed, baseline modifications already appear to exist in *Lis1* mutant developing brains and no further additive effects are generated by *TBC1D3* expression. It is likely that early Lis1 depletion by 50%, while not inducing heavily deleterious effects, generates sufficient cellular modifications to prevent TBC1D3 from generating bRG-like cells. Thus, Lis1 is necessary for murine bRG-like cell formation associated with expression of a hominoid-specific gene contributing to bRG generation in humans.

## Results

### TBC1D3 promotes the generation of bRG-like cells in the mouse

The bRG population is naturally scarce in the developing mouse brain, we therefore expressed *TBC1D3* to amplify this population (Ju et al., 2016). To test this tool under our experimental conditions, WT mice were first electroporated at E14.5 with pCS2-cMyc-*TBC1D3* (Ju et al, 2016) together with a pCAG-IRES-tdTomato plasmid to identify electroporated cells, compared to pCAG-IRES-tdTomato alone (Figure 1A-B’). Mice were sacrificed two days later. TBC1D3 expression led to the presence of tdTomato-positive (tdTomato+) basally-positioned cells with somata localized in the SVZ and IZ (Figure 1B’). These possessed either a basal process or both an apical and basal process, as described previously for bRG cells (Betizeau et al., 2013). Immunohistochemistry showed that many basal cells with bRG-like morphology were Pax6+ (Figure 1B’, B’’). Quantifications showed a significant increase of the proportion of Pax6+ cells among electroporated cells when TBC1D3 is expressed compared to control. This includes when considering the entire cortical wall (Total, 10.9% ± 1.9 for control versus (vs.) 24.5% ± 0.1 for TBC1D3, p<0.01), as well as when dividing the cortical wall into the VZ (36.9% ± 3.5 for control vs. 49.9% ± 7.7 for TBC1D3, p<0.01), SVZ (5.5% ± 0.9 for control vs. 15.1% ± 4.5 for TBC1D3, p<0.05) and IZ (1.9% ± 0.5 for control vs. 11.8% ± 2.6 for TBC1D3, p<0.05) (Figure 1C). This suggests that TBC1D3 expression increases numbers of Pax6+ RG which leave the VZ and move towards more basal regions.

**Figure 1:**
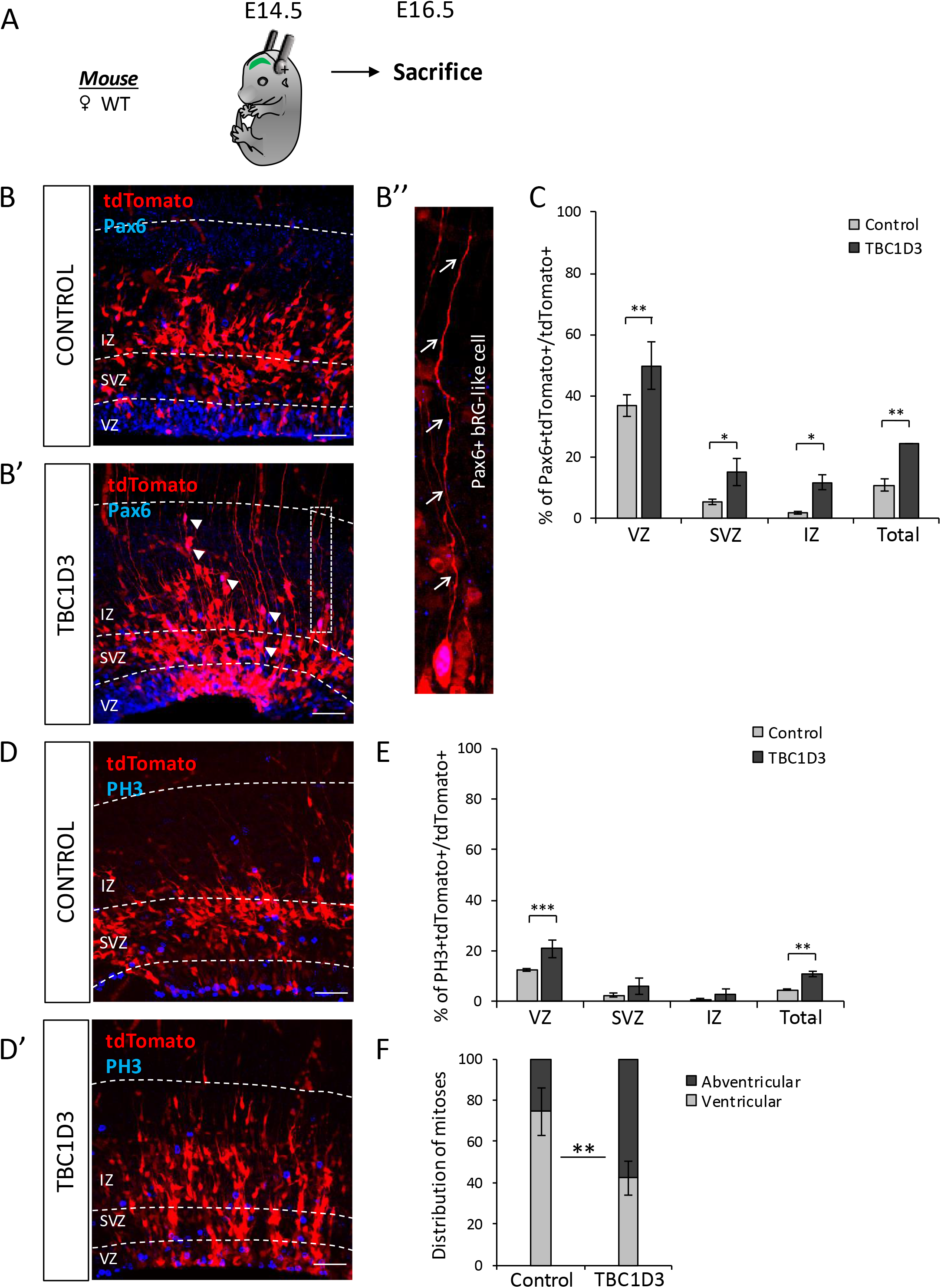
TBC1D3 expression leads to the amplification of Pax6+ cells and the generation of bRG-like cells. **A)** Schematic view of the protocol used for experiments in Figure 1: WT embryos (Swiss genetic background) were electroporated at E14.5 and sacrificed at E16.5. **B,B’,B’’)** Pax6 immunohistochemistry (blue) of E16.5 electroporated mouse brains. Control (tdTomato, B) and TBC1D3 (tdTomato + TBC1D3, B’) (arrowheads point to Pax6+ bRG-like cells in different regions of the cortical wall). Enlargement shows a Pax6+ bRG-like cell with typical morphology and showing a long basal process (arrows, B’’). **C)** Proportion of Pax6+ tdTomato+ / tdTomato+ cells in each cortical region and in the whole cortex (‘Total’) per ROI in control (grey bars) or TBC1D3 (black bars) electroporated brains (3 x 50 μm slices per brain, 3 animals per condition, 2-way ANOVA with Bonferoni *post hoc*, * : p<0,05, ** : p<0,01). **D,D’)** PH3 immunohistochemistry (blue) of E16.5 electroporated mouse brains. Control (tdTomato, D) and TBC1D3 (tdTomato + TBC1D3, D’). **E)** Proportion of PH3+ tdTomato+ / tdTomato+ in each cortical region and in the whole cortex (‘Total’) per ROI in Control or TBC1D3 electroporated brains (3 x 50 μm slices per brain, 3 animals per condition, 2-way ANOVA with Bonferoni *post hoc*, ** : p<0,01, *** : p<0.001). Note the overall increase of mitoses in TBC1D3 electroporated brains. **F)** Distribution of ventricular mitoses versus abventricular mitoses (3 x 50 μm slices per brain, 3 animals per condition, *t*-test, ** : p<0.01). Note the fact that the majority of mitoses take place at the ventricular surface in the control condition while TBC1D3 brains display a shift towards more abventricular mitoses. Scale bars (B,B’,D,D’) = 50 μm.

The effect of TBC1D3 expression on mitoses was assessed by performing phospho-Histone 3 (PH3) immunostaining (Figure 1D, D’). TBC1D3 expression led to increased numbers of PH3+ mitotic cells compared to the control plasmid. This was significant when considering the proportion of mitotic cells amongst fluorescent cells across the whole cortical wall (4.5% ± 0.2 for control vs. 10.8% ± 1.1 for TBC1D3, p<0.01), as well as specifically when assessing the VZ (12.6% ± 0.5 for control vs. 20.8% ± 3.4 for TBC1D3, p<0.001) (Figure 1E) and tendencies were also observed in the SVZ (2.38% ± 0.81 for control vs. 6.13% ± 3.16 for TBC1D3) and IZ (0.81% ± 0.52 for control vs. 2.50% ± 2.4 for TBC1D3). aRG mitoses generally take place at the ventricular surface in the control condition (Figure 1D), related to interkinetic nuclear migration (Kulikova et al., 2011; Taverna et al., 2014). When TBC1D3 was expressed, the location of mitoses displayed a notable basal shift. Indeed, the proportion of abventricular mitoses increased (Total: 43.0% ± 11.0 for control vs. 69.1% ± 1.5 for TBC1D3, p=0.052, data not shown), with a decreased proportion of divisions at the ventricular surface. Considering only abventricular mitoses in the VZ (Figure 1F), there were 25.6% ± 11.2 for control vs. 57.5% ± 8.2 for TBC1D3, p<0.01) (Figure 1F). The latter suggests that basal mitoses were more likely to occur in the presence of TBC1D3. Overall, these results are consistent with the original study (Ju et al, 2016) suggesting that TBC1D3 expression in mouse brain progenitors promotes the generation of bRG-like cells.

### Forebrain-specific *Lis*1 knockout severely perturbs cortical development

After validating the production of TBC1D3-induced bRG-like cells, *Lis1* mutants were generated. Various Cre lines have been used in the past to induce *Lis1*^fl/fl^ recombination, showing defects depending on the timing of Lis1 depletion, targeted cell types and the dosage of the Lis1 remaining (Gambello et al., 2003; Hirotsune et al., 1998; Yingling et al., 2008). We set out to deplete *Lis1* in a forebrain-specific manner at an early stage, before the onset of neurogenesis, to ensure depletion of this gene in all neural progenitors. We used a new *Lis1*^fl/fl^ stock from which the neo-cassette had been removed (referred to here as *Lis1*^fl/fl^). Indeed, the presence of the neo-cassette was previously shown to alter *Lis*1 gene expression (Gambello et al., 2003; Hirotsune et al., 1998), generating *de facto* an hypomorphic allele even in the absence of the Cre recombinase. To deplete *Lis1* in neo-removed *Lis1*^fl/fl^ mice, an Emx1-IRES-Cre knockin mouse was used (referred to here as Emx1-Cre) to induce recombination in neural progenitors starting at E9.5 (Gorski et al., 2002). To verify the pattern of expression of the Cre, Emx1-Cre animals were crossed with the Rosa26-EGFP^RCE/RCE^ line (Sousa et al., 2009). Brains of the embryos showed GFP expression restricted to the cortex, as well as in fiber tracts including the corpus callosum and the anterior commissure (data not shown).

After having confirmed the recombination pattern in the Emx1-Cre mouse line, *Lis1*^fl/fl^ animals were crossed with Emx1-Cre animals, to obtain Lis1^fl/+^ Emx1-Cre^+/−^ mutant mice. These were then crossed with Lis1^fl/fl^ mice to obtain WT (*Lis1*^fl/+^ or *Lis1*^fl/fl^; Emx1-Cre^+/+^), heterozygote (HET, *Lis1*^fl/+^; Emx1-Cre^+/−^) and knockout (KO, *Lis1*^fl/fl^; Emx1-Cre^+/−^) animals in the same litter. Postnatal day 0 (P0) brains of HET pups did not show any macroscopic cortical differences when compared to control, consistent with previous studies (Gambello et al., 2003; Hirotsune et al., 1998). However, brains of KO animals were largely devoid of a cortex (Figure 2A, C). This phenotype was easily visible at E14.5 (data not shown), and KO animals died between P3 and P7. This result further confirmed that reducing the dosage of *Lis*1 by half does not have obvious major deleterious effects on mouse cortical development (and HETs survive normally), whereas further decrease has severe consequences (Gambello et al., 2003; Hirotsune et al., 1998).

**Figure 2:**
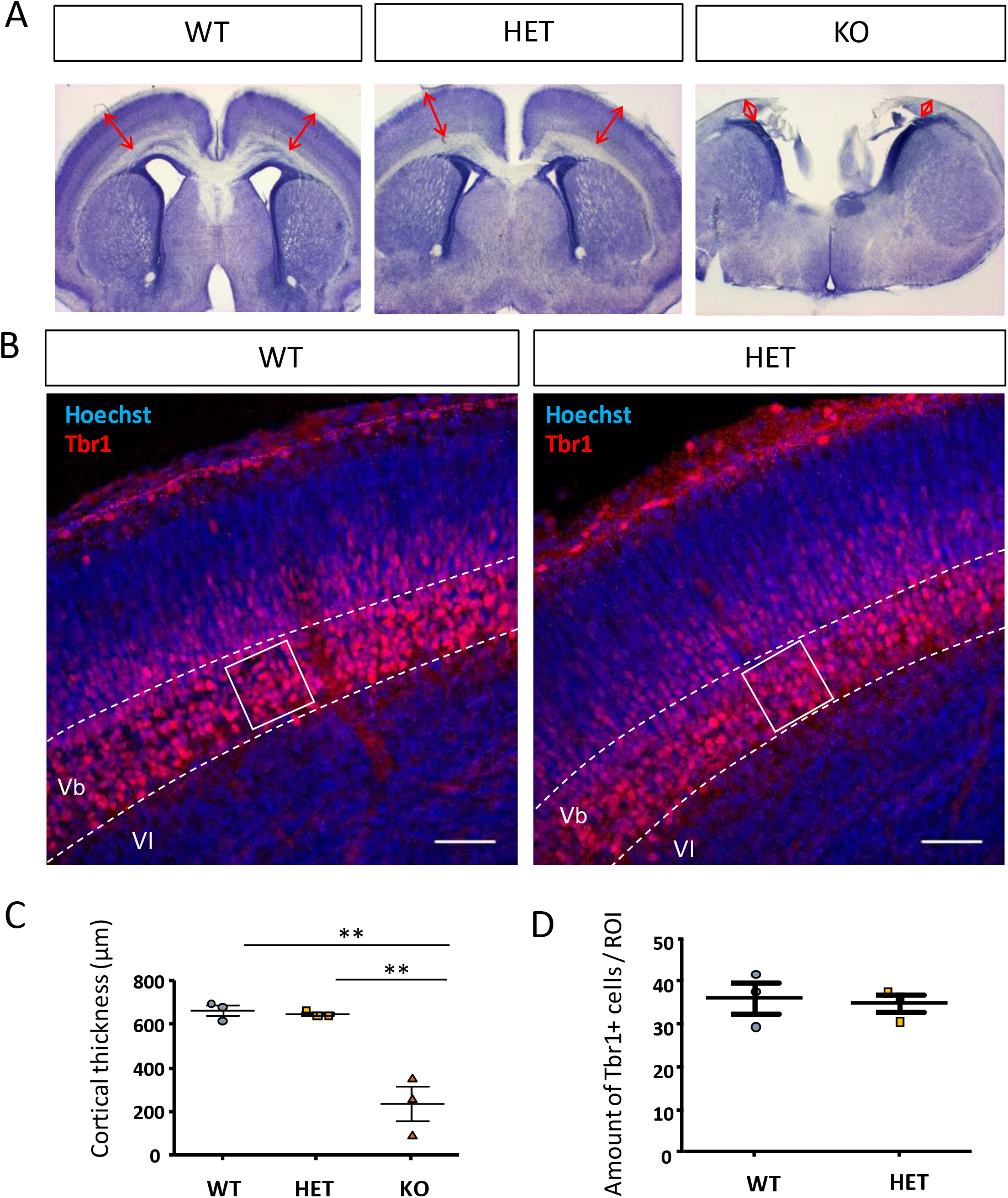
Lis1 inactivation in forebrain neural progenitors severely impairs neocortical development in the KO, while heterozygotes show no major effects. **A)** Nissl staining of Lis1 WT, HET and KO P0 brain slices. HET brains show no macroscopic differences compared to WT animals. The cortex and medial areas in KO brains appear severely affected (absence of cortex and corpus callosum). **B)** Tbr1 immunohistochemistry (red) at E16.5 in WT and HET mouse brains to label deep-layer neurons. Nuclei counterstained with Hoechst dye. White squares display typical ROIs used for quantification of cell number. **C)** Quantification of cortical thickness (indicated by red arrows in A, 3 animals per condition, one-way ANOVA, ** : p<0.01) showing no differences between WT and HET mice, and a large reduction in KO mice.. **D)** Quantification of the number of Tbr1 cells per ROI (3 x 50 μm slices per brain, 3 animals per condition, *t*-Test). No significant difference was observed between the genotypes.

Considering that the cortex is almost totally absent in KO animals, making a fine assessment of Lis1 function in KO progenitors less appropriate, we continued our experiments comparing only WT and HET animals. To that end, *Lis1*^fl/fl^ mice were crossed with Emx1-Cre^+/−^ animals to obtain WT (Cre negative) or HET embryos in the same litter. To evaluate whether Lis1 depletion by 50% had an impact on cortical organization, even though no major defects were observed macroscopically, Tbr1 immunostaining was performed. No significant differences were observed between WT and HET mice when assessing the number of Tbr1+ deep layer neurons (Figure 2 B, D). This together with the cortical thickness measurements suggests that in the HET state, cortical neuron production and migration occur relatively normally.

Performing the same crosses and generating pregnant females, embryos were then electroporated at E14.5 with pCS2-cMyc-TBC1D3 or control (pCS2-cMyc) plasmid together with the pCAG-IRES-tdTOMATO reporter plasmid, and mice were sacrificed 2 days later. Activated caspase 3 (aCas3) immunostaining was first performed to assess cell death. No significant differences were observed, when depleting Lis1, or expressing TBC1D3 (Supplementary Figure 1A, B).

Overall, these results with early forebrain inactivation of *Lis1* showed that full KO at E9.5 leads to severe cortical developmental defects, while depletion of *Lis1* by 50% has no obvious effect on cortical layering organization and a non-significant impact on cell death.

### Lis1 depletion prevents bRG-like cell amplification upon TBC1D3 expression

We next investigated whether, in the presence of TBC1D3, Lis1 depletion would alter or prevent the generation of bRG-like cells. To evaluate the effect of TBC1D3 (or the control plasmid) expression on RGs in the *Lis1* mutant cortex, Pax6 and Ki67 co-immunostaining was performed (Figure 3B-E’). Pax6+ tdTomato+ cells were assessed across the cortical wall and in the different zones (Figure 3F). Under these conditions, no significant differences in the percentages of Pax6+ cells within the tdTomato population were observed in the VZ and SVZ regions. The overall percentage of Pax6+ cells amongst the tdTomato+ cells across the cortical wall also did not show significant differences, although there was a tendency for increase in the TBC1D3 WT condition. In the IZ, no differences were noted between WT and HET brains electroporated with the control plasmid, however a significant increase in the proportion of Pax6+ tdTomato+ cells was observed when TBC1D3 is expressed in WT brains (9.0% ± 3.0 for Control WT vs. 33.2% ± 9.4 for TBC1D3 WT, p<0.05). However, when TBC1D3 is expressed in HET brains, numbers of Pax6+ cells and levels are similar in the IZ to that of the HET condition electroporated with the control plasmid (7.8 ± 6.5 for Control HET vs. 13.3% ± 3.0 for TBC1D3 HET, no significant differences). This suggests that Lis1 depletion largely prevents bRG-like cell amplification in the presence of TBC1D3.

**Figure 3:**
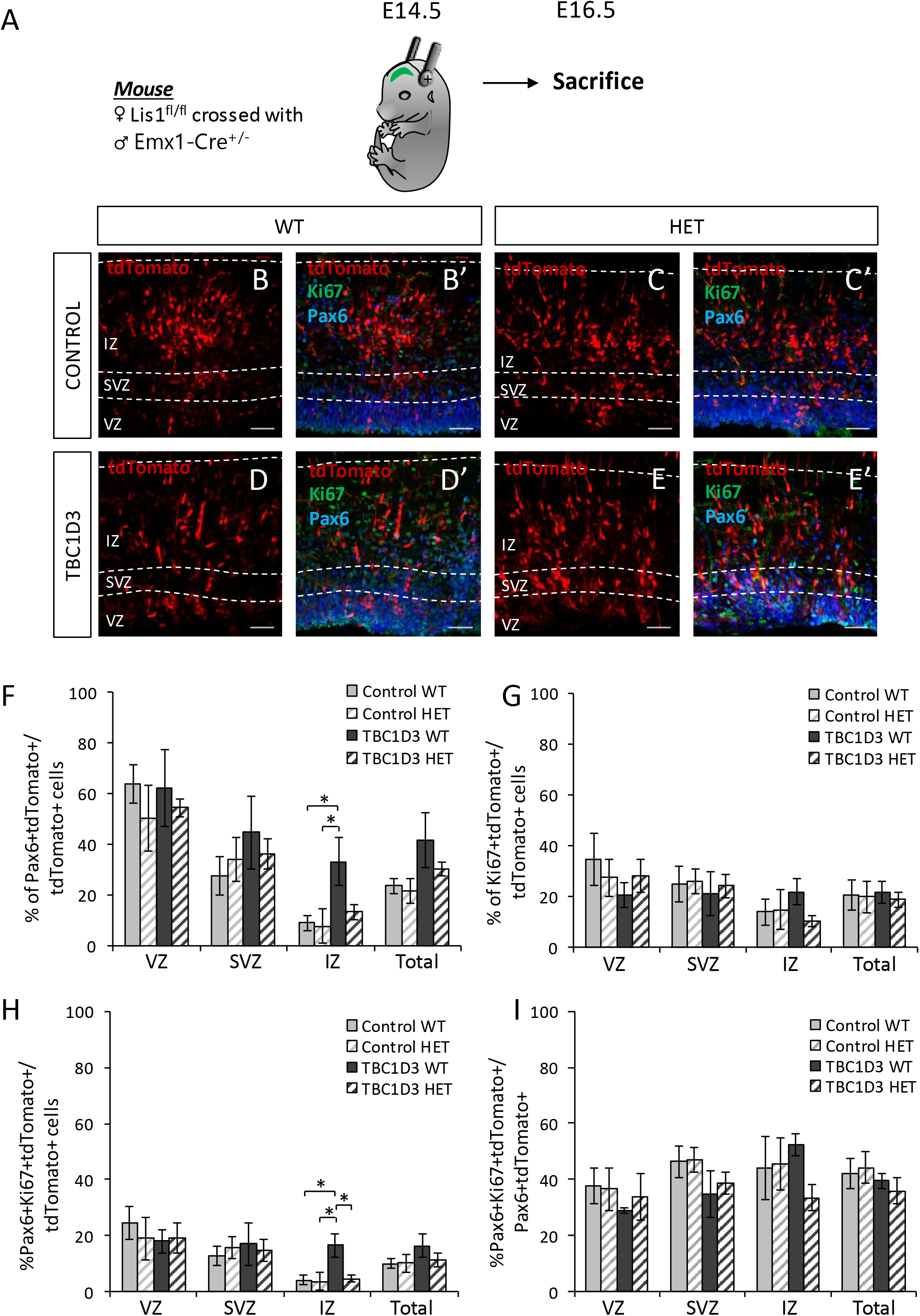
*Lis*1 depletion does not allow the generation of cycling bRG-like cells in the IZ in the presence of TBC1D3. **A)** Schematic view of the protocol used: embryos from *Lis1*^fl/fl^ females crossed with Emx1-Cre^+/−^ males were electroporated at E14.5 and sacrificed at E16.5. **B – E’)** Pax6 (blue) and Ki67 (green) immunohistochemistry in E16.5 WT (B, B’, D, D’) and HET (C, C’, E, E’) mouse brains after electroporation of pCS2-cMyc-Control + tdTomato (Control, B, B’, C, C’) and pCS2-cMyc-TBC1D3 + tdTomato (TBC1D3, D, D’, E, E’) plasmids. tdTomato alone and the merge of tdTomato, Pax6 and Ki67 staining are shown for each condition. **F, G, H, I)** Proportion of Pax6+ tdTomato+ / tdTomato+ (F), Ki67+ tdTomato+ / tdTomato+ (G), Pax6+ Ki67+ tdTomato+ / tdTomato+ (H) and Pax6+ Ki67+ tdTomato / Pax6+ tdTomato+ (I) cells in WT and HET brains electroporated with control or TBC1D3 plasmids. Results are presented as percentages across the cortical wall (‘Total’) or within each cortical region (3 x 50 μm slices per brain, n=4 animals per condition, 2 Way ANOVA with Tukey *post hoc*, *: p<0,05). Note TBC1D3 significantly increases Pax6+ and Pax6+Ki67+ proportions in the IZ, and this increase is prevented by Lis1 depletion. Scale bars (B-E’) = 50 μm.

The effect of Lis1 depletion on cycling RG was then assessed by quantifying the proportion of cells co-labeled with Pax6 and Ki67, a marker of proliferation (Figure 3G, H). Overall proportions of Ki67+ cells did not differ significantly between the genotypes and conditions (Figure 3G). However, when considering Pax6+ Ki67+ tdTomato+ triple labeled cells (Figure 3H), similar results to those described in Figure 3F were obtained: in the IZ no difference was found between WT and HET conditions with the control plasmid, however TBC1D3 expression in WT animals resulted in an increased proportion of Pax6+ Ki67+ cells (4.2% ± 1.5 for Control WT vs. 16.7% ± 4.1 for TBC1D3 WT, p<0.05). This increase was not observed with Lis1 depletion (4.7% ± 1.2 for TBC1D3 HET vs. 16.7% ± 4.1 for TBC1D3 WT vs, p<0.05). This suggests that TBC1D3 expression in WT induces an increased production of cycling bRG-like Pax6+ cells in the IZ, and importantly, Lis1 depletion prevents this phenomenon from occurring.

To assess whether Lis1 dosage also influences the ability of Pax6+ RGs to cycle, we measured the proportion of cycling RG (Pax6+ Ki67+ tdTomato+) among electroporated RG (Pax6+ tdTomato+) (Figure 3I). No significant differences were observed, suggesting proliferation of RGs was not greatly affected.

Overall, these combined results suggest that TBC1D3 expression is no longer able to promote the generation of bRG-like cells and cycling bRG-like cells in the IZ when Lis1 has previously been depleted by 50%.

### Lis1 depletion with TBC1D3 expression does not alter Tbr2+ cell numbers

We then investigated whether the population of Tbr2+ IPs was also affected in Emx1-Cre Lis1 mutants. Tbr2 and Ki67 co-immunostainings were performed (Figure 4A-D’) and cells were quantified. There were notably no significant differences when considering either the total or the regional numbers of electroporated Tbr2+ IPs (Figure 4E), Tbr2+ Ki67+ cycling IPs (Figure 4F) or the ability of Tbr2+ IPs to cycle (Figure 4G), including comparing TBC1D3 WT and TBC1D3 HET conditions. These results suggest that only RG progenitors are affected by Lis1 depletion when TBC1D3 is expressed.

**Figure 4:**
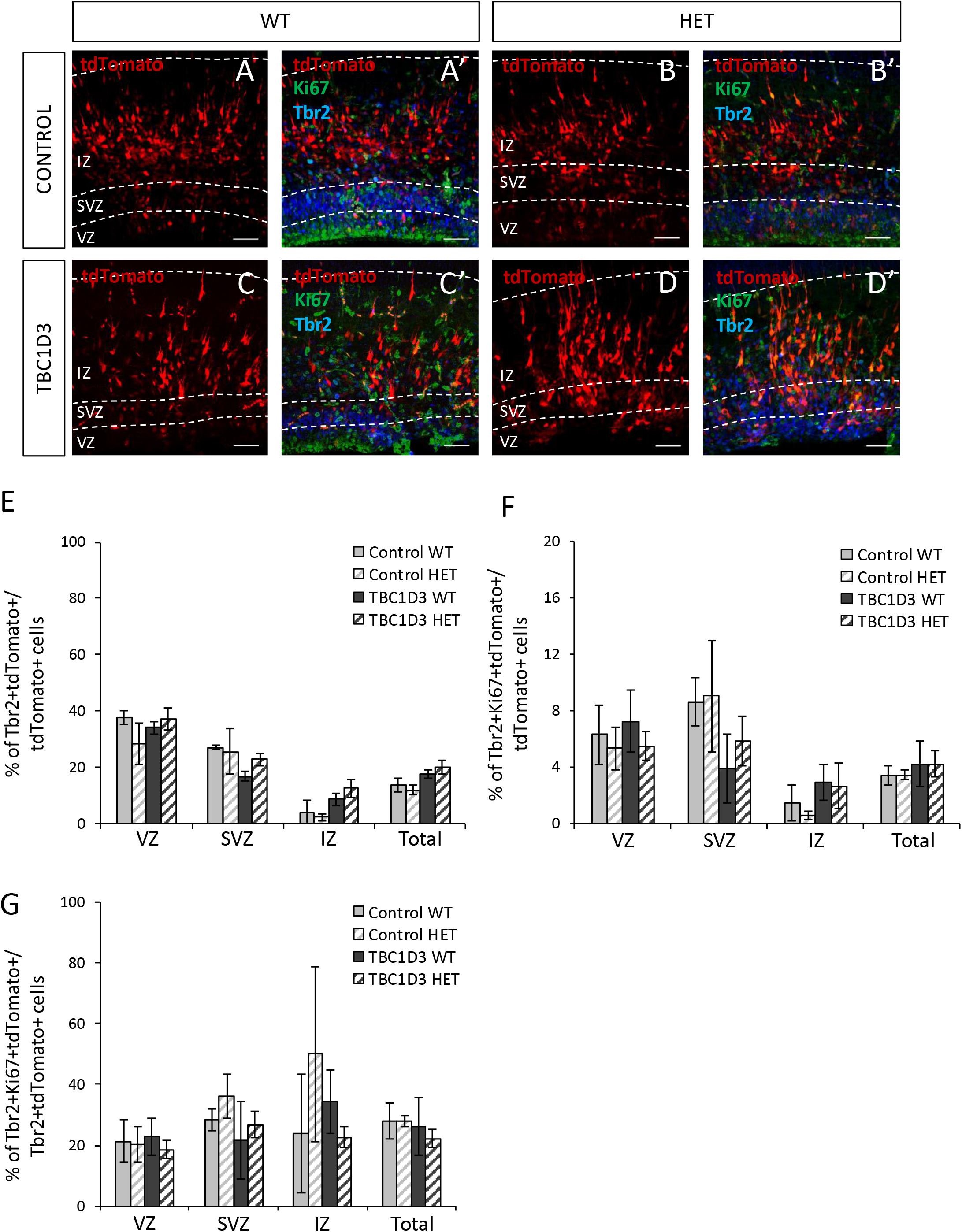
Lis1 depletion has no effect on intermediate progenitors upon TBC1D3 overexpression. **A) A – D’)** Tbr2 (blue) and Ki67 (green) immunohistochemistry in E16.5 WT (A, A’, C, C’) and HET (B, B’, D, D’) mouse brains after electroporation at E14.5 of pCS2-cMyc-Control + tdTomato (Control, A, A’, B, B’) and pCS2-cMyc-TBC1D3 + tdTomato (TBC1D3, C, C’, D, D’) plasmids. tdTomato alone and the merge of tdTomato, Ki67 and Ki67 staining are shown. **E, F, G)** Proportion of Tbr2+ tdTomato / tdTomato+ (E), Tbr2+ Ki67+ tdTomato+ / tdTomato+ (F), Tbr2+ Ki67+ tdTomato+ / Tbr2+ tdTomato+ (G) in WT and HET brains electroporated with control or TBC1D3 plasmids. Results are represented as percentages across the cortical wall (‘Total’) or within each cortical region (3 x 50 μm slices per brain, n=5-8 animals per condition for Tbr2+ tdTomato+ / tdTomato analysis, n= 3 animals per condition for other analyses, 2 Way ANOVA with Tukey *post hoc*). Scale bars (A-D’) = 50 μm.

### N-Cadherin expression is perturbed in Lis1 deficient mice but not upon TBC1D3 expression

To assess how Lis1 depletion may inhibit the generation of bRG-like cells, we first investigated whether VZ organization and cell interactions were affected. Indeed, expression of TBC1D3 in the mouse was previously shown to decrease the expression of N-Cadherin, and this alteration may explain how bRG-like cells are generated potentially by delamination in this model (Ju et al., 2016). To test if *Lis1* haploinsufficiency impairs this mechanism, N-Cadherin staining was performed followed by immunofluorescence intensity quantification at the ventricular surface. No obvious effect of TBC1D3 expression on N-Cadherin was observed in WT animals, contrary to the original study. However, when Lis1 was depleted, both with control and TBC1D3 plasmids, decreased expression of N-Cadherin was observed (Figure 5A).

**Figure 5:**
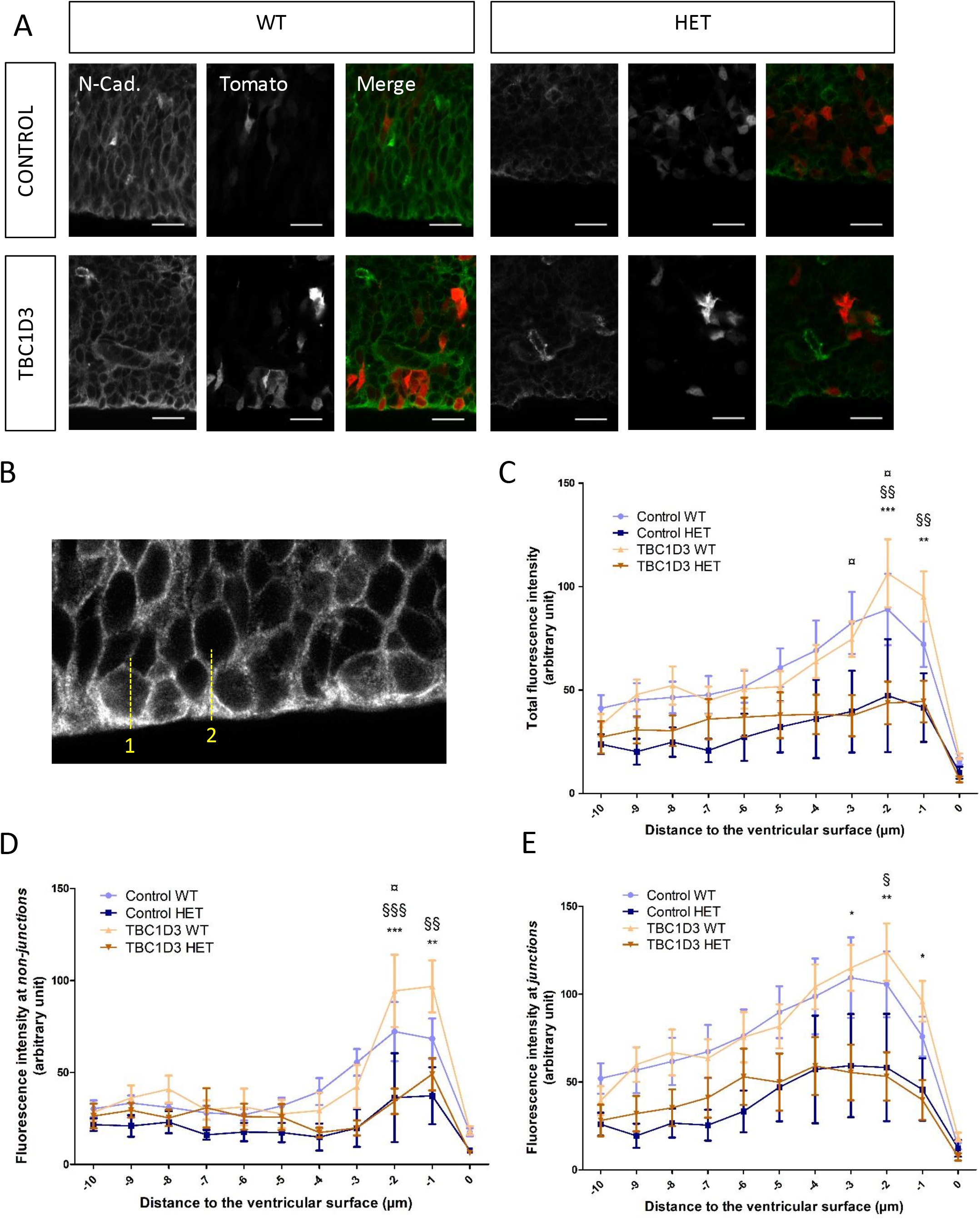
Lis1 depletion perturbs N-Cadherin expression at the ventricular surface. **A)** N-Cadherin immunohistochemistry in E16.5 WT and HET mouse brains after electroporation at E14.5 with pCS2-cMyc-Control + tdTomato (Control) and pCS2-cMyc-TBC1D3 + tdTomato (TBC1D3) plasmids. Figure shows N-Cadherin (N-Cad), tdTomato (Tom), and merge panels for each condition. **B)** Quantification procedure for N-Cadherin fluorescence: 10 μm lines contacting perpendicularly to the ventricular surface were drawn on ImageJ, and fluorescence intensity was measured every micron along the lines. Quantification was performed at non-junctions sites (meaning, in the middle of cells, line 1), at the cell adhesion sites (meaning in between cells, line 2), and for the total by combining both. **C, D, E)** Quantification of N-Cadherin total (C), “*non-junction*” (D) and “*junction*” (E) fluorescence intensity (arbitrary unit) at the ventricular surface every μm along a 10 μm line perpendicular to the ventricular surface. The extremity of the ventricular surface is marked as 0 on the X axis. (3 x 50 μm slices per brain, 3-4 animals per condition, 2-way ANOVA for repeated measures with Bonferoni *post hoc*, *, **, *** : p<0,05, p<0.01, p<0.001 respectively between TBC1D3 WT and TBC1D3 HET, §, §§, §§§ : p<0,05, p<0.01, p<0.001 respectively between Control HET and TBC1D3 WT, ¤ : p<0.05 between Control WT and TBC1D3 HET). Scale bars (A) = 20 μm.

Performing quantifications across electroporated areas, in control WT animals, a peak of fluorescence intensity was obvious close to the ventricular surface where N-Cadherin expression is the strongest (Figure 5B). Two-way Anova for repeated measures showed a significant interaction for fluorescence intensity between the conditions and distance from the ventricle (p < 0.0001). No significant differences were observed between Control WT and TBC1D3 WT, suggesting TBC1D3 expression does not affect N-Cadherin expression. Comparing WT to Het conditions however, we noticed a tendency for a decrease comparing Control HET to Control WT, and a significant decrease comparing TBC1D3 HET to TBC1D3 WT (*) close to the ventricular surface (Figure 5B). This suggests that Lis1 depletion reduces N-Cadherin expression at the ventricular surface.

To assess more precisely the effect of Lis1 depletion and TBC1D3 expression on N-Cadherin expression at the ventricular surface, we performed measurements outside of cell junctions (“*non-junctions*”) and at cells junctions (“*junctions*”) (Figure 5C, D). Overall, results were similar to those described above, i.e. Control HET and TBC1D3 HET animals showed lower intensity of fluorescence. At the *non-junction*, interaction between conditions and distance from the ventricle was significant (p = 0.0002) and significant differences were observed between TBC1D3 WT and TBC1D3 HET (*), between TBC1D3 WT and Control HET (§) and between Control WT and TBC1D3 HET (¤), in regions close to the ventricular surface. At *junctions*, the interaction was also significant (p=0.0011), and the overall tendency of HETs having lower intensities as compared to WT equivalent brains was observed, with TBC1D3 HET showing a significantly lower intensity of fluorescence as compared to TBC1D3 WT. Overall, these results suggest that Lis1 depletion alters N-Cadherin expression, potentially leading to perturbed function of VZ progenitors.

### Both Lis1 depletion and TBC1D3 expression alter ventricular mitoses and spindle orientations

We then investigated whether the Emx1-Cre *Lis1* mutation altered other aspects of VZ progenitor cell function, helping to prevent TBC1D3 expression from inducing the production of bRG-like cells. To evaluate whether mitosis was affected, PH3 immunostaining was performed and the proportion of PH3+ mitotic cells quantified (Figure 6A). No significant differences were observed in the overall and regional proportions of mitotic electroporated cells between the different conditions and genotypes (Figure 6B). However, when considering the distribution of mitoses in the VZ, we observed that Lis1 HET depletion (after electroporation with the control plasmid) promoted significantly more basal abventricular mitoses compared to WT (27.6% ± 9.1 for Control WT vs. 64.6% ± 6.6 for Control HET, p<0.05, Figure 6C), consistent with previous *Lis*1 mutation studies (Yingling et al., 2008). TBC1D3 expression in WT animals also promoted an increased proportion of basal abventricular mitoses (27.6% ± 9.1 for Control WT vs. 62.8% ± 5.9 for TBC1D3 WT, p<0.05), consistent with the original study (Ju et al., 2016). Interestingly, when TBC1D3 was expressed in HET animals, the proportion of abventricular mitoses was similar to levels of HET mice electroporated with the control plasmid, or WT with the TBC1D3 plasmid (60.0% ± 9.9 for TBC1D3 HET vs. 64.6% ± 6.6 for Control HET). Thus, both conditions (Lis1 HET control and WT TBC1D3) lead to increased proportions of abventricular mitoses individually, and expression of TBC1D3 combined with Lis1 depletion does not show potentiation of the effect. Since Lis1 HET aRG mitoses are already perturbed when TBC1D3 is introduced, this may help to explain why bRG-like cells are not generated in the mutant model.

**Figure 6:**
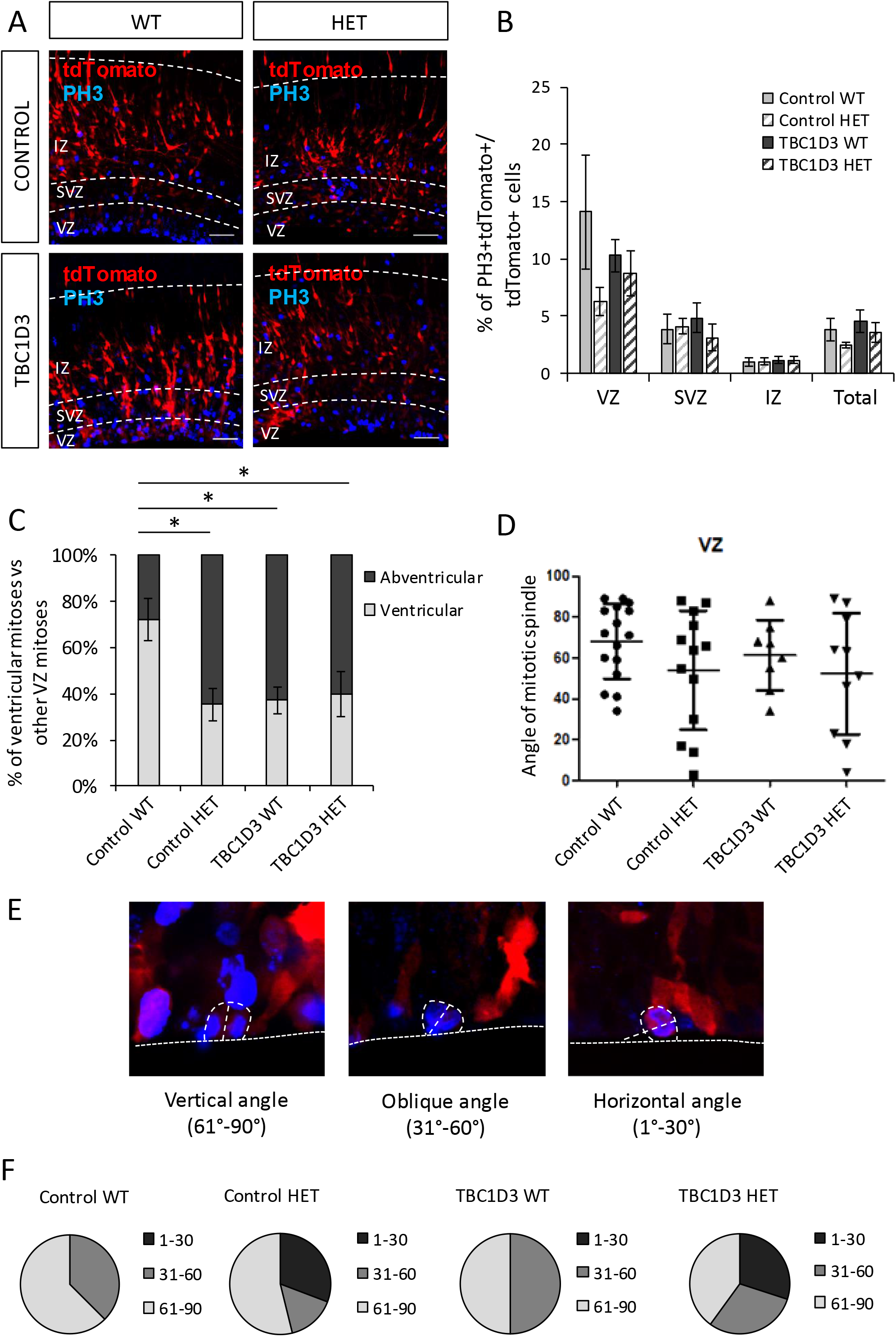
Both Lis1 and TBC1D3 increase abventricular mitoses but only Lis1 perturbs mitotic spindle orientations. **A)** PH3 immunohistochemistry (blue) in E16.5 WT and HET mouse brains after electroporation at E14.5 with pCS2-cMyc-Control + tdTomato (Control) and pCS2-cMyc-TBC1D3 + tdTomato (TBC1D3) plasmids. **B)** Proportion of PH3+ tdTomato+ / tdTomato+ in each cortical region and in the whole cortex (‘Total’) per ROI in WT or HET mice electroporated with Control or TBC1D3 plasmids (3 x 50 μm slices per brain, 7 animals per condition, 2-way ANOVA with Bonferoni *post hoc*). **C**) Distribution of ventricular mitoses versus abventricular mistoses (3 x 50 μm slices per brain, 7 animals per condition, 2-way ANOVA with Bonferoni *post hoc*, * : p>0,05). Note that the majority of mitoses take place at the ventricular surface in the control WT condition, while control HET, TBC1D3 WT and TBC1D3 HET brains display a significant shift towards abventricular mitoses. **D)** Distribution of mitotic spindle angles in electroporated cells dividing at the ventricular surface. Note wider variability for Lis1 HET conditions with resulting lower angle mean. **E)** Mitotic spindle angles (categorized in vertical (90°-61°), oblique (60°-31°) and horizontal (30°-1°)) in electroporated cells dividing at the ventricular surface. Note that *Lis*1 HET depletion dysregulates mitotic spindle angles by generating horizontal angles at the expense of vertical and oblique both in control and TBC1D3 conditions. For D,E 8-16 cells in 3 brain slices of n=6-8 animals per condition. F) Example of aRG mitoses for which the mitotic spindle angles were measured. Scale bar (A) = 50 μm.

We next decided to assess mitotic spindle orientations in *Lis1* mutants in the presence or absence of TBC1D3. The angles of division of fluorescent dividing aRGs, in contact with the ventricular surface and engaged in anaphase, were measured (Figure 6D,E, F). Angle distributions for WT and HET with control plasmids were akin to what was shown previously (Yingling et al 2008). When Lis1 is depleted by 50%, a lower angle mean was observed, with an apparently wider variability (Control WT: mean = 68.1°, standard deviation = 18.36; Control HET: mean = 54.0°, standard deviation = 29.16, Levene test: p=0.068). TBC1D3 expression in WT did not appear to greatly alter mitotic spindle angles compared to the control plasmid, however when TBC1D3 was expressed in *Lis1* mutant brains, we observed the same tendency as with the control plasmid, Lis1 depletion induced a lower angle mean and a wider variance (Figure 5D, TBC1D3 WT: mean = 61.4°, standard deviation = 17.17; TBC1D3 HET: mean = 52.4°, standard deviation = 31.36, Levene test: p=0.10).

The naturally low numbers of electroporated cells performing mitosis at the moment of the sacrifice and fixation of the embryos prevented further meaningful statistical analyses related to the variability of the angles. Hence, to provide further insight into the data, angles were clustered in three groups (61°-90° for horizontal divisions, 31°-60° for oblique divisions and 1°-30° for vertical divisions, Figure 5E). Lis1 depletion showed more variable angle values, notably increased values were observed between 1°-30° (0 % in WTs versus 30 % in HETs), apparently at the expense of 31-60° angles. Expression of TBC1D3 in WT increased oblique divisions (31-60°, 50 % with TBC1D3 versus 37.5 % with the Control plasmid). TBC1D3 expression in Lis1 HET brains appeared to randomize angles (with values observed of 30 %, 30 % and 40% for 1°-30°, 31°-60° and 61°-90° respectively, Figure 5F).

Thus, early alteration of mitotic spindle angles in *Lis1* HET animals could prevent proper segregation of cell fate determinants at the time of TBC1D3-induced divisions, which might explain how Lis1 prevents bRG-like cell enrichment in the presence of TBC1D3.

## Discussion

Basal RGs were identified relatively recently and shown to be important for the expansion of the neocortex in gyrencephalic species (Betizeau et al., 2013; Florio et al., 2015; Hansen et al., 2010; Fietz et al., 2010; Reillo et al., 2011). However, few studies to-date associate defects in these cells with cortical malformations, most probably since these cells are few in the rodent, often used to model these neurodevelopmental disorders. In order to overcome these limitations, we used genetic tools to amplify these cells in the mouse, assessing their production in *Lis1* mouse mutants. Using a newly developed *Lis*1 allele, we show that forebrain-specific conditional KO leads to a severe cortical defect and death in early postnatal stages. Taking advantage of viable heterozygote mice (mimicking the gene dosage in human patients), we tested the production of bRGs in Lis1 mutant conditions. We show that bRG-like cells do not form correctly when TBC1D3 is expressed in *Lis1* HET mutants. We demonstrate that *Lis1* mutants show severely perturbed spindle orientations and reduced N-Cadherin expression from early cortical development which appears to preclude the production of bRG-like cells. We did not observe N-Cadherin impairments in TBC1D3 WT animals, differing from a previous study (Ju et al., 2016). This difference could be due to genetic background, since Ju and colleagues used C57Bl6 mice, while in our study, *LIS1*^fl/+^ *Emx1-Cre*^+/−^ animals had a hybrid FVB/C57Bl6 background.

Different Cre-expressing lines have been used previously to deplete *Lis1* (Yingling et al., 2008). Use of Pax2-Cre with Cre expression starting E8 (Rowitch et al., 1999), expected to impact neuroepithelial cells, leads to a severe impairment of neuroepithelium function in KO animals, with midbrain and hindbrain degeneration and highly increased apoptosis in forebrain cells. Similar to Emx1-Cre KOs studied here, pups did not survive after birth. Using GFAP-Cre mice with Cre expression starting at E12.5 and expected to impact aRGs (Brenner and Messing, 1996) there is a slight decrease of the cortical thickness in KOs, affecting both deep and superficial neurons, and an almost absent hippocampus. With the Emx1-Cre mouse line, we depleted Lis1 at an intermediary timepoint (E9.5), at the beginning of the transition from neuroepithelial cells to aRGs in the forebrain (Gorski et al., 2002). We observed an almost complete degeneration of the cortical wall in KOs, akin to the Pax2-Cre phenotype and much stronger than that of GFAP-Cre. This suggests that Lis1 is critical for the proper transition from neuroepithelial cells to aRGs, the start of neurogenesis and the correct acquisition of cell fate.

LIS1 heterozygote mutations in human lead to a spectrum of disorders with varying grades of severity, ranging from pachygyria with a rostro-caudal gradient to complete agyria, subcortical band heterotopia and/or microcephaly (Leventer et al., 2001; Lo Nigro et al., 1997; McInnes et al., 2007; Romero et al., 2018). Thus, haploinsufficient (or heterozygous) mutations have severe consequences in human (Di Donato et al., 2017), while depletion by 50% in mice shows few defects of cortical development (Gambello et al., 2003; Hirotsune et al., 1998; Yingling et al., 2008). In *Lis1* HET animals in our study, no significant changes were observed in the production or cycling capabilities of RGs or IPs. Neuron production and cortical layering also appeared grossly unaffected, consistent with previous studies (Gambello et al., 2003; Pramparo et al., 2010). Thus *Lis1* HET mutation in the mouse from E9.5 only leads to comparatively mild defects, suggesting either the activation of compensatory factors (Pawlisz et al., 2008; Pramparo et al., 2010), or that in human, cortical development involves mechanisms not present in the mouse. Indeed, mice are naturally lissencephalic, while the human cortex is gyrencephalic and much more complex (Borrell, 2019).

TBC1D3 is a hominoid specific gene, present in one copy in the chimpanzee and 8 in human (Hodzic et al., 2006). By *TBC1D3* overexpression in the mouse, it is possible to “humanize” the mouse cortex, enriching it in primate-like cells, and therefore potentially sensitizing the mouse cortex to mutations that have deleterious effects in humans (Ju et al., 2016). Interestingly, when TBC1D3 is overexpressed, HET and WT mice display differences, notably in numbers of bRG-like cells. This might reveal how bRGs could contribute to the human phenotype. Interestingly, it has already been shown that bRGs have perturbed mitosis in human MDS organoids (Bershteyn et al., 2017), exhibiting a contiguous deletion of 17p13 including LIS1. Our experiments in the *Lis1* mutant mouse differ since bRG production in the VZ was tested due to this single gene mutation. It is important to continue testing the role of Lis1/LIS1 affecting the production and amplification of this cell type.

We hence focused on bRG-like cell production at mid-corticogenesis in *Lis1* HET mouse mutants. bRG producing mechanisms have been associated with either alterations of mitosis spindle orientation leading to more oblique and vertical divisions (Kalebic et al., 2018; Liu et al., 2017; Wang et al., 2016; Wong et al., 2015), and/or to detachment from the apical surface via decreased cell adhesion (Ju et al., 2016; Martínez-Martínez et al., 2016; Narayanan et al., 2018; Tavano et al., 2018; Taverna et al., 2014). Many of the genes that were described to increase bRG-like cell generation induce more oblique divisions (Penisson et al., 2019), and mitotic spindle orientation is a well-described mechanism that may contribute to the fine tuning of aRG daughter cell fate (LaMonica et al., 2013; Taverna et al., 2014). Lis1 could contribute to this phenomenon by regulating dynein-dynactin complex activity (Coquelle et al., 2002; Htet et al., 2020; Wang et al., 2013), as well as actomyosin-mediated cell membrane contractility, influencing cleavage plane positioning (Moon et al., 2020). However, the mechanisms by which TBC1D3 promotes the expansion of the bRG-like cell pool in the mouse remain poorly understood. TBC1D3 expression was shown previously to reduce the expression of N-Cadherin at the ventricular surface, with concomitant decreased expression of Trnp1 and activation of the MAPK pathway (Ju et al., 2016; Stahl et al., 2013). However, the exact mechanisms initiated by these modifications remain unclear (Penisson et al., 2019) and notably we did not observe reduced N-Cadherin in our experimental conditions. It is also unclear how the N-Cadherin defects already present in *Lis1* mutant cells influence these processes to prevent the generation of bRG-like cells.

N-Cadherin and adherens junctions in general are known to maintain proper organization of the neuroepithelium (Stocker and Chenn, 2015). In organoids derived from MDS patient cells, N-Cadherin expression was also altered in the VZ and this phenotype was rescued with LIS1 re-expression (Iefremova et al., 2017). This suggests a further role for Lis1, confirmed by our results, in the maintenance of apical junctions and the overall structure of the ventricular surface, which may be critical for bRG production.

Both *Lis1* depletion and *TBC1D3* expression led to an increased proportion of abventricular mitoses. These might explain progenitor detachment, in fitting with the increased numbers of bRG-like cells when TBC1D3 is expressed. However, this phenomenon is not sufficient to explain the cellular identity of abventricularly dividing cells in *Lis1* mutants. During interkinetic nuclear migration (Kulikova et al., 2011; Tavano et al., 2018), mitoses take place at the ventricular surface. Dynein is required for apical migration towards the ventricle (Tsai et al., 2010) and this process may be perturbed in Emx1-Cre *Lis1* mutants, leading to ectopically dividing ‘aRGs’. If apical migration is slower, or severely inhibited, this would increase the number of mitoses away from the ventricular surface. Such an ectopic position of the nucleus might be expected to impair *TBC1D3*-inducing mechanisms to generate bRG-like cells, since in our model, *TBC1D3* expression was induced in aRGs, after deletion of Lis1. Thus, we do not exclude that ectopic Pax6+ mitoses contribute to impairing bRG-like production, however, further defects were also observed at the ventricular surface in *Lis1* mutants.

The effect of Lis1’s depletion on mitotic spindle orientation has been well characterized both in mice and organoids. When Lis1 expression is decreased in early apical progenitors (neuroepithelial or aRGs), performing mainly horizontal divisions (with vertical cleavage planes), they start to engage in oblique and vertical divisions (Yingling et al., 2008; Pramparo et al., 2010; Iefremova et al., 2017). Furthermore, LIS1 expression in MDS organoids rescued the mitotic spindle angle phenotype (Iefremova et al., 2017), showing the importance of this gene for this phenotype. In our study, when comparing Control WT to Control HET, and TBC1D3 WT to TBC1D3 HET, we observed an angle dispersion increase, associated with *Lis1* mutation, as well as a small increase in the proportion of oblique divisions in presence of TBC1D3. Importantly though, the latter was quite mild compared to the *Lis1* phenotype and furthermore, the same angle of division could lead to different outcomes, i.e. oblique divisions in *Lis1* mutants may not engage daughter cells in the same fate as *TBC1D3* expression. It is possible that under *Lis1* mutant conditions, TBC1D3 may not be able to induce a proper segregation of cell fate determinants and / or the maintenance of the basal process, required for the epithelial fate (Moon et al., 2020). Regulation of mitotic spindle orientation and mitosis progression is managed in part by a Lis1-Ndel1-dynein complex (Moon et al., 2014). Similar to the stabilization of microtubules, re-expression of Lis1 or overexpression of Ndel or dynein led to rescue of spindle misorientation in *Lis1* mutant MEFs (Moon et al., 2014). We hence predict that an early rescue of spindle orientation in *Lis1* mutant aRGs, or a different timing of *Lis1* mutation, might enable TBC1D3 to induce bRG-like cell production.

Our work combined with the previous MDS organoid study (Bershteyn et al., 2017) begin to decipher bRG mechanisms in *Lis1/MDS* mutant conditions. In our mouse mutant work, only TBC1D3 is experimentally manipulated without the possibility of multiple entries into the bRG pathway. Alternatively, human cells in developing organoids are likely to express a larger battery of genes and signaling pathways involved in the generation of bRGs (Penisson et al., 2019). However, as the state of bRGs in human pathologies remains understudied, using a single gene e.g. TBC1D3, in mouse mutants can be useful for assessing specific pathways and helping us to further understand bRG mechanisms. bRGs have been suggested to be critical for the correct development of the cortex in gyrencephalic species, due to their number and proliferative properties. However, whether their number, distribution or function are perturbed in pathologies in which cortical folding is altered, i.e. in the human lissencephaly brain, instead of organoids or mouse models, still remains unknown. Investigating human-like bRG cells induced in mouse models, produced with tools that can even lead to the generation of mouse cortical folds, can open new windows to study and revisit MCDs.

## Methods

### Animals

Research was carried out conforming to national and international directives (directive CE 2010/63 / EU, French national APAFIS n° 8121) with protocols followed and approved by the local ethical committee (Charles Darwin, Paris, France). Mice were housed with a light/dark cycle of 12 h (lights on at 07:00).

Swiss wild-type (WT) mice (Janvier Labs, France) were used for *in utero* electroporation to confirm the TBC1D3 effect on bRG-like cell production (Ju et al., 2016). For the study of Lis1 depletion, Lis1 floxed allele mice (Hirotsune et al., 1998) maintained on the FVB background from which the neo selection cassette had been removed (obtained from Osaka University) were crossed with C57Bl6J Emx1-Cre knockin animals (Gorski et al., 2002). The Rosa26-EGFP^RCE/RCE^ mouse line was used as reporter (Sousa et al., 2009). All mice were housed in the IFM institute animal facility or at the CDTA, Orléans, France. Genotyping of all transgenic mouse lines was performed via PCR using these oligonucleotides:

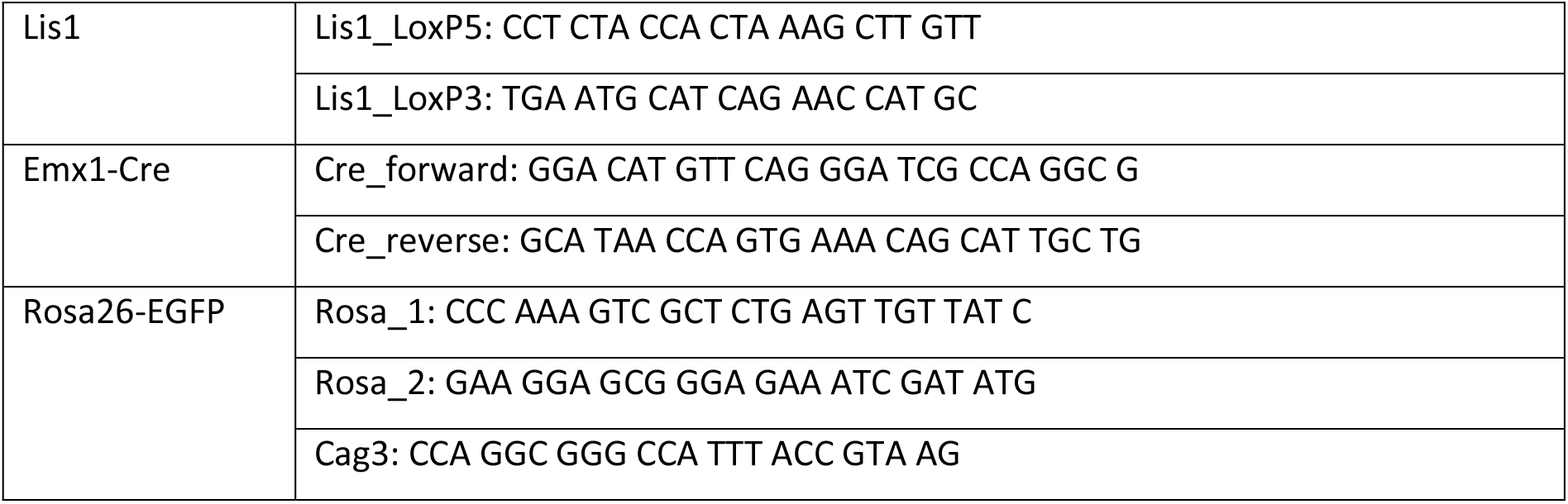

### Plasmid preparation

Plasmids were amplified from NEB 5α bacterial cultures transformed with pCS2-cMyc-TBC1D3 (gratefully received from Zhen-Ge Luo’s team), pCS2-cMYC-Control (generated from the TBC1D3 construct) or pCAG-IRES-TdTomato plasmids (kindly provided by C. Lebrand, Lausanne), all of which carried the ampicillin resistance gene. These were cultured overnight (O/N) on petri dishes containing culture medium and ampicillin (50 mg/L). Clones were then amplified and cultured in 200 mL LB medium + ampicillin (50 mg/L) O/N. Plasmid DNAs were then extracted using the NucleoBond™ Xtra midi EF kit (Fisher Scientific), following the recommended protocol. After extraction, plasmid preparations were resuspended in endonuclease-free H2O at approximately 5 μg/μL, concentrations verified with spectrophotometry (NanoDrop 1000 spectrophotometer, Thermo Scientific) and stored at −20°C.

### Electroporation in utero

E14.5 pregnant females were anaesthetized with 4 % isoflurane (Baxter) and a Minerve Dräger Vapor 2000 apparatus, and maintained on a heating pad at 37°C at 2-3% isoflurane depending on animal weight and respiratory rhythm. The abdomen of the animals was sectioned and uterine horns were extracted and kept hydrated with NaCl 0.9% (Braun) + penicillin streptomycin (P/S, 10 mg/mL). A plasmid solution (DNA concentration of 1 μg/μl + Fast-Green 1% (F-7258, Sigma)) was then injected in the lateral ventricles of the embryos with glass micropipettes. Micropipettes were generated from glass capillaries (GC150TF-10, Harvard Apparatus) that were pulled using a Narishige PC-100 pipette puller. Electrodes (CUY650P3, NepaGene) were placed on the sides of the head of embryos in such a way as to propagate entry of the plasmid in the pallium. Electroporation was performed with 5 pulses of 35 V, each with a duration of 50 ms, and with 950 ms intervals using a CUY21 (NepaGene) electroporator. The uterus was then replaced carefully inside the mother’s abdomen, the inside of the abdomen was washed with NaCl 0.9% + P/S solution and the wound was closed with stitches using Vycril thread (VICRYL JV390, ETHICON) for the muscle wall, and, Ethilon thread (ETHILON F3206, ETHICON) for the skin. Mice were injected subcutaneously with Flunixin (4 mg/kg) and put back in their cage on a heating pad at 37 °C to recuperate with monitoring.

### Brain slicing, histology, immunofluorescence and confocal acquisition

Females that underwent *in utero* electroporation were sacrificed 48 hours later, and brains of embryos were collected and fixed in 4 % paraformaldehyde overnight at 4 °C. Brains were then rinsed and conserved in 1X PBS (diluted from PBS 10X, ET330 Euromedex) + 0.1 % azide (S200-2, Sigma-Aldrich). Prior to Vibratome sectioning, brains were embedded in an 8 % sucrose (200-301-B, Euromedex), 6 % agarose (LE-8200-B, Euromedex) PBS solution. Serial 50 μm thick coronal sections were performed with a VT1000S vibratome (Leica). For immunofluorescence labelling, 2 to 3 slices per brain were used for every experiment. Saturation and permeabilization was performed for 1 hour with a 10 % normal goat serum (NGS) (16210072, Gibco) and Triton X-100 (T9284, Sigma) solution in 1X PBS, then slices were incubated with primary antibodies O/N (or for 72 hours for Ki67) diluted in permeabilization and saturation buffer at 4 °C under agitation. Primary antibodies include Pax6 (Biolegend, BLE901301, 1:300), PH3 (Millipore, 06-570, 1:400), Tbr2 (AB23345, Abcam, 1:300), Tbr1 (grateful gift from Robert Hevner, 1:1000), aCas3 (559565, BD Pharmigen, 1:250), Ki67 (BD Biosciences, 556003, 1:300), N-Cadherin (BD Biosciences, 610920, 1:500). Ki67 antibody required antigen retrieval: prior to permeabilization and saturation, slices were incubated in 10 mM sodium citrate, 0.05 % Tween pH 6.0 at 95 °C for 20 minutes, then left to cool down at room temperature before rinsing 3 times with 1X PBS. After incubation with primary antibodies, slices were rinsed 3 times with permeabilisation and saturation buffer. Secondary antibodies were then incubated at 1:800 dilution for 2 hours at room temperature under agitation, the list include donkey-anti-rabbit Alexa 488 (ThermoFisher, A21206), donkey-anti-mouse Alexa 488 (ThermoFisher, A21201), donkey-anti-rabbit Alexa 647 (ThermoFisher A31573). Slices were then rinsed with Hoechst (33258, Sigma) 1:10000 for 10 minutes diluted in 1X PBS, and then rinsed twice with 1X PBS. Slices were mounted on Superfrost Plus microscopic slides (Thermo Scientific) with Fluoromount-G (Invitrogen). Confocal acquisitions were performed with an SP-5 Leica microscope (x40), stacks of 30-40 μm were performed.

### Histology

For studies in mouse pups (P0), animals were rapidly decapitated, and brains were removed and postfixed in 4% paraformaldehyde overnight at 4°C and then stored in PBS. Then, dissected brains were embedded in 6% agarose and 8% sucrose. 50-μm coronal sections were prepared using a vibratome (VT1000S; Leica), Nissl staining was performed on sections mounted on Superfrost slides (Thermo Fisher). The sections were analyzed with a brightfield microscope (Provis; Olympus) using a charge-coupled device (CCD) camera (CoolSNAP CF; Photometrics) and 4× (NA = 0.13) objective.

### Analysis

Quantification for Tomato, Pax6, Sox2, Tbr1, Tbr2, Ki67, aCas3 and PH3, as well as mitotic spindle angles was performed using Icy Bioimage software. Regions of interest (VZ, SVZ, IZ, CP) were defined in actual, adjacent or similar level sections using Hoechst, Pax6, Sox2 and/or Tbr2 staining. VZ was defined using Pax6 staining, the IZ was characterized as a region of lower cell density using Hoechst, and the SVZ was defined either by Tbr2 staining or as the region lying in between predefined VZ and IZ. Cells positively stained for tdTomato and other nuclear markers, or aCas3 clusters, were counted with the “Manual Counting” plug-in according to their localization. Results are shown as a percentage of tdTomato cells, to account for variabilities in efficiency of *in utero* electroporation. Quantification for N-Cadherin fluorescence intensity was performed using Fiji software. Ten 20 μm lines were drawn perpendicular to and across the ventricular surface such that the middle of each line (10^th^ μm) was at the transition between the ventricle and the ventricular surface, 5 lines were drawn at the junction between 2 cells, and 5 not at the junction (i.e. in the middle of cells). Multi plotting for pixel intensity was performed with ROI manager for each line. Values for measurements within the ventricles were discarded because ImageJ generated artefactual random values between 1 and 10. Values were also simplified for statistical analysis, one value of fluorescence per μm was kept out of the 163 values for 20 μm that ImageJ generated. The means of the 5 lines for the junctions, 5 lines for the non-junctions and 10 in total per image were then calculated. Cortical thickness was measured using Fiji software. 3 lines per hemisphere was drawn, and mean was calculated. Statistical analysis was performed using R and the Rcmdr graphical interface, and Prism software (GraphPad). Normality of residuals and homoscedasticity was checked with Shapiro and Levene tests respectively prior to performing any parametrical test (*t*-tests, one-way and two-way ANOVAs).

---

**Supplementary Figure 1. Lis1 depletion in the heterozygous state with TBC1D3 expression does not lead to increased cell death.** A) Activated Caspase 3 immunohistochemistry (aCas3, blue) in E16.5 WT and HET mice after IUE at E14.5 with tdTomato + pCS2-cMyc-Control (Control) and tdTomato + pCS2-cMyc-TBC1D3 (TBC1D3) electroporated brains. **B)** Quantification of aCas3 clusters per ROI in Control WT and HET, and TBC1D3 WT and HET animals. No significant differences were observed, but there were tendencies for an increase when Lis1 is depleted and for when TBC1D3 is expressed. Scale bars (C,G) = 50 μm.

